# Gustatory sensitivity to amino acids in bumblebees

**DOI:** 10.1101/2024.11.28.625904

**Authors:** Sergio Rossoni, Rachel H. Parkinson, Jeremy E. Niven, Elizabeth Nicholls

**Author notes:** **Authors for correspondence:** Sergio Rossoni, Elizabeth Nicholls.

## Abstract

Bees rely on amino acids from nectar and pollen for essential physiological functions. While nectar typically contains low (<1 mM) amino acid concentrations, while their levels in pollen are higher, but vary widely (10-200 mM). Behavioural studies suggest bumblebees have preferences for specific amino acids but whether such preferences are mediated via gustatory mechanisms remains unclear. This study explores bumblebees’ (*Bombus terrestris*) gustatory sensitivity to two essential amino acids (EAAs), valine and lysine, using electrophysiological recordings from gustatory sensilla on their mouthparts. Valine elicited a concentration-dependent response from 0.1 mM, indicating that bumblebees could perceive valine at concentrations found naturally in nectar and pollen. In contrast, lysine failed to evoke a response across tested concentrations (0.1-500 mM). The absence of lysine detection raises questions about the specificity and diversity of amino acid-sensitive receptors in bumblebees. Bees responded to valine at lower concentrations than sucrose, suggesting comparatively higher sensitivity (EC_50_: 0.7 mM *vs*. 3.91 mM for sucrose). Our findings indicate that bumblebees can rapidly evaluate the amino acid content of pollen and nectar using pre-ingestive cues, rather than relying on post-ingestive cues or feedback from their nestmates. Such sensory capabilities likely impact foraging strategies, with implications for plant-bee interactions and pollination.

## 1. Introduction

Bees’ ability to exploit floral resources efficiently depends upon their evaluation of the food rewards offered. Bees base their nectar-based foraging decisions on parameters including sugar concentration [1], flow rate [2] and distance from the nest [3]. In addition to carbohydrates obtained from nectar, bees also require protein for maintenance, sexual maturation and larval development [4]. Amino acids are the building blocks of proteins and can be obtained from both pollen and nectar [5]. While highly variable between species and even individual plants [6,7], amino acids are typically found in pollen at 10-200 mM [8–11] and, although found at much lower concentrations in nectar (< 1 mM), are still the most abundant nutrient after sugars [12–15].

Amino acids influence bee feeding and foraging choices in compound- and concentration-dependent ways [16–18]. Bees prefer to collect pollen with higher levels of amino acids [19], though some amino acids can increase nectar consumption at low concentrations but inhibit consumption at high concentrations *e*.*g*. proline [20,21]. Bees also prefer nectars containing one or more essential amino acids (EAA) that cannot be synthesised and must be obtained from the diet [22], over those containing only non-essential amino acids (NEAA) [15,23].

Whether the effects of amino acids on bees’ dietary choices are mediated by pre-ingestive taste cues, remains uncertain [4,15]. The apparent regulation of amino acid intake by bees at the individual and colony level could arise from post-ingestive processes or, in the case of social bees, feedback from nestmates or larvae. A behavioural study using chemo-tactile conditioning of the proboscis extension response (PER) [24] found that when presented to their antennae, bumblebees can distinguish some EAAs (*e*.*g*. lysine) and NEAAs (*e*.*g*. cysteine) from water but not others (e.g. valine and proline, respectively). Moreover, bees could not discriminate between different amino acids [24]. However, PER assays are unable to disentangle olfactory, gustatory and tactile cues; honeybees can smell certain amino acids at high concentrations, albeit beyond those found naturally in nectar [25]. The honeybee gustatory receptor *AmGr10* binds to several amino acids when expressed in cell lines, and electrophysiological recordings show gustatory sensilla on honeybee mouthparts (galea) respond to two NEAAs, glutamate and aspartate [26].

Here, we conducted the first electrophysiological recordings of bees’ gustatory responses to two EAAs, valine and lysine, to test whether bumblebees’ (*Bombus terrestris*) can detect amino acids pre-ingestively via gustatory sensilla. According to behavioural evidence, lysine is reported to be perceived by bumblebees, while valine is not [24], and both were tested across a range of concentrations (0.1-500 mM) encompassing amino acid concentrations found in both nectar and pollen [11,12].

## 2. Materials and methods

### (a) Animal husbandry and selection

Bumblebee (*Bombus terrestris audax*) colonies were obtained from Biobest (Westerlo, Belgium) (*n*=2) and Koppert (Berkel en Rodenrijs, The Netherlands) (*n*=1). Colonies were maintained at the University of Sussex, U.K., housed either within a flight cage (73×73×65 cm) or connected to a feeding arena (40×40×35 cm) via a corridor (4×4×26 cm). Bees had *ad libitum* access to a nectar substitute (Biogluc, Biobest, Westerlo, Belgium) via both ground and suspended feeders, and finely ground honeybee collected pollen (Agralan, Swindon, U.K.), presented on chenille stems placed inside white plastic cups. Nectar was replenished as needed and pollen changed daily. Workers observed collecting pollen from the feeders were marked on the thorax with a small dot of white enamel paint (Humbrol, Hornby Hobbies Limited, Margate, U.K.). Only individuals observed collecting pollen at least once were used in testing (*n*=27).

### (b) Animal preparation and restraint

Bumblebees were caught in the feeding arena and chill-immobilised at 4 °C overnight. On the day of recording, bees were inserted in small plastic tubes and harnessed using small strips of Parafilm M (American National Can, Greenwich, CT, U.S.A.). Bees were then transferred to a plate of sealing wax beneath a microscope (Nikon AZ100, Tokyo, Japan). Using wax, the head, antennae, and front legs of the bumblebee were immobilised, and the glossa and labial palps were manually extended and immobilised to the plastic tube. The galeae were rinsed in ultrapure water and dried with QL100 filter paper (Fisherbrand, Fisher Scientific, Loughborough, U.K.), and subsequently extended and fixed to the plastic tube using small strips of Parafilm to prevent heat damage.

### (c) Electrophysiological recordings

Extracellular tip recordings (figure 1*a*) were performed on A-type sensilla chaetica on the left galea [27]. A 25 μm tungsten wire (Alfa Aesar, Ward Hill, MA, U.S.A.) was inserted into the galea at the proximal end and pushed gently down to ∼1 mm from the sensilla from which the recording was to be made. This wire was used as the reference electrode. The recording electrode was a 250 μm silver/silver chloride wire, placed inside a borosilicate glass capillary (1×0.58×100 mm (ODxIDxL); Harvard apparatus, Holliston, MA, U.S.A.) pulled on a P-97 micropipette puller (Sutter Instrument Co., Novato, CA, U.S.A.) to a tip diameter of ∼20-50 μm to fit comfortably over A-type gustatory sensilla on the galea (*n*=124) without deflecting the hair. Capillaries were filled with tastant solutions (see below) with no added electrolytes. The capillary was moved close to the galea using an LBM-7 manipulator (Scientifica, Uckfield, U.K.) and contact with the apical pore was made using an MO-203 micromanipulator (Narishige, Tokyo, Japan), mounted on the LBM-7.

**Figure 1.**
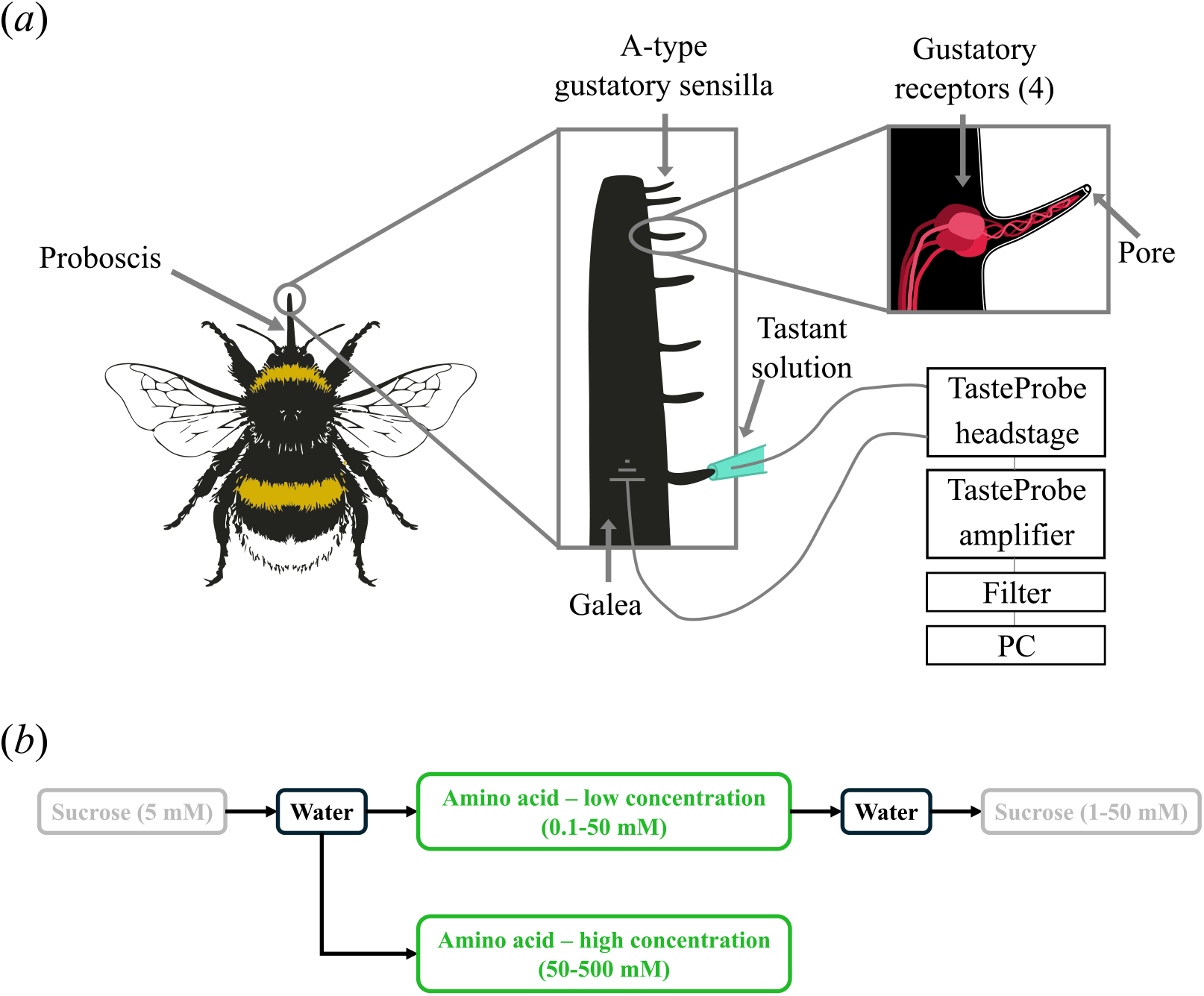
(*a*) The experimental set-up for recording from gustatory sensilla on the bumblebee galea. (*b*) The stimulation protocol.

Electrical signals, which represent the ensemble activity of the four gustatory receptor neurons (GRNs) innervating each sensillum (figure 1*a*), were recorded through the tastant solutions. The electrodes were connected to a TasteProbe [28] headstage and TasteProbe DTP-02 10x amplifier (Syntech, Buchenbach, Germany). Recordings were made for 20 s with a 1 Hz high-pass filter. The signal was further amplified with 5x gain and band-pass filtered between 10 and 10k Hz (LHBF-48X filter-amplifier, npi electronic GmbH, Tamm, Germany). Recordings were digitised via a CED micro1401-3 digital-to-analogue converter (Cambridge Electronic Design Ltd, Cambridge, U.K.) and acquired using CED Spike2 software (10.09) at 20k Hz.

### (d) Tastant stimulation and protocol

Gustatory sensilla on the galeae (*n*=3-6 sensilla per bee) were stimulated with aqueous solutions of D-(+)-sucrose (Fisher Scientific, Loughborough, U.K.), L-valine (Thermo Fisher Scientific, Heysham, U.K.), or L-lysine (Thermo Fisher Scientific, Waltham, MA, U.S.A.). Syringes filled with tastant solutions were prepared in advance and stored at -10 °C. Prior to the electrophysiological experiments, the syringes were thawed, and stored at 4 °C. The capillary used for stimulation was filled immediately prior to use, to minimise water evaporation affecting solution concentration. Bees were allocated to one of two experimental groups, one stimulated with low amino acid concentrations and one with high concentrations (figure 1*b*), to prevent slow adaptation over the course of the experiment. In both conditions, gustatory sensilla were tested initially with 5 mM sucrose and water, to ensure that sucrose sensitivity was comparable between GRN ensembles in all conditions. Bees in the low-concentration group were exposed to increasing concentrations of either lysine or valine from 0.1 to 50 mM. Then, bees in this group were tested with increasing concentrations of sucrose from 1 to 50 mM [27]. The high-concentration group was exposed to increasing concentrations of either lysine or valine from 50 to 500 mM. At least 3 minutes elapsed between exposure to different tastants and concentrations to reduce possible adaptation. Temperature was monitored every 30 s throughout the recording, using an automated temperature logger (EasyLog-USB-2-LCD, Lascar Electronics, Whiteparish, U.K.).

### (e) Data processing

Individual recordings were imported into Matlab (R2024a, MathWorks inc., Natick, MA, U.S.A.) for offline analysis based on [29]. Recordings were trimmed either using contact artefacts, or by the 20 s timestamp acquired from the amplifier, whichever was shortest; the entire trimmed length was used for data extraction. Filtered recordings were then created using a band-pass second-order Butterworth filter between 100 and 1000 Hz. Using these copies, a threshold was then selected manually for each recording to identify spikes. The peak of the spikes above the threshold was used to acquire 4 ms waveforms from the raw signal. The waveforms in each recording were inspected. Four measures were used to distinguish spikes and artefacts: maximum voltage reached, minimum voltage reached, waveform amplitude, and waveform half-width. Individual waveforms with unusual or irregular shapes were considered movement artefacts and removed. Spike frequency was calculated as the total number of spikes in the recording divided by the duration after trimming. To describe the change in spike frequency to increasing stimulation of tastants, we fitted a sigmoid using the method of non-linear least squares in the ‘stats’ package using R studio (4.4.1 R software [30]). For the final figures, electrophysiological recordings were smoothed with a 15 Hz high-pass Butterworth filter and taken from 20 ms after trimming.

### (e) Statistical analysis

Statistical tests were implemented using R studio software, version 2024.04.2, and R software, version 4.4.1 [30]. Homogeneity of variance was tested using Levene’s test in the ‘car’ package. Two datasets with Gaussian sampling distribution and homogeneous variance were compared using a Student’s *t*-test in the ‘stats’ package. A generalised linear mixed effects model (GLMM) implemented in the ‘lmerTest’ package was used to test whether tastant concentrations affected spike frequency: log_10_(spike frequency)∼log_10_[amino acid]+(1|temperature)+(1|sensillum). A GLMM was also used to compare the concentration dependency between valine and sucrose: log_10_(spike frequency)∼substance*log_10_[tastant]+(1|temperature)+(1|sensillum). To avoid mathematical infinity when using transformed data at 0 mM or 0 Hz, 1 was added to all spike frequencies and tastant concentrations prior to logarithmic transformation. Where relevant, GLMMs with significant predictors were followed by *post hoc* pairwise comparisons with Hommel’s adjusted *p-*values in the ‘stats’ package.

## 3. Results

We observed reliable activation of the gustatory receptor neurons (GRNs) in response to sucrose (figure 2*a*). Higher sucrose concentrations (>25 mM) typically triggered burst-like activity in the GRNs (figure 2*a*) [27]. GRNs also responded to stimulation with the amino acid valine between 0.1 and 500 mM (figure 2*b*), though no burst-like activity was recorded during valine stimulation, even at high (500 mM) concentrations. In contrast, little activity was seen during stimulation with the amino acid lysine, even at high (500 mM) concentrations (figure 2*c*).

**Figure 2.**
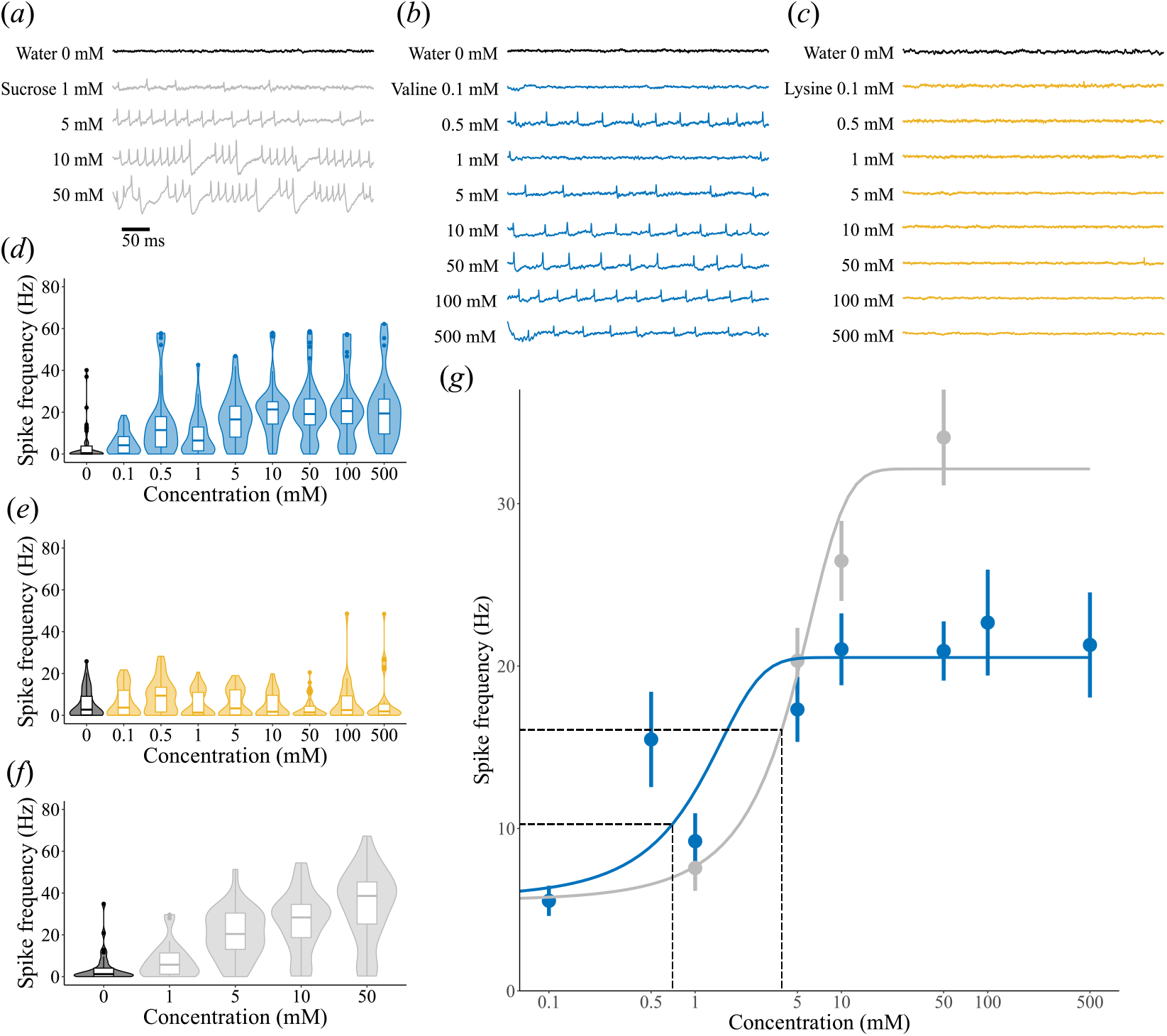
Gustatory neurons in sensilla show concentration dependency in response to valine and sucrose but not to lysine. (*a*) Example recording of one gustatory sensillum in response to water (black) and increasing concentrations of sucrose (grey). (*b*) As in *a* but for valine (blue). (*c*) As in *a* but for lysine (yellow). (*d*) The spike rate (Hz; *med* and *IQR*) of gustatory neurons over ∼20 s in response to water (black) and increasing concentrations of valine (blue). (*e*) As in *d* but for lysine (yellow). (*f*) As in *d* but for sucrose (grey). (*g*) Sigmoid fits to the spike rates for valine (blue) and sucrose (grey). Dashed lines indicate the half maximal effective concentration (EC_50_) and its corresponding spike frequency.

Valine concentration was a significant predictor of GRN spike frequency (*n*=59, GLMM: t=13.7, coefficient estimate 0.349, standard error (se) 0.026, *p*<0.001), indicating that sensitivity to valine was concentration dependent (figure 2*d*). While stimulation with 0.1 mM valine did not significantly increase the spike frequency above baseline (*n*=93, pairwise comparison post-hoc, *p*=0.054), 0.5 mM valine significantly increased the spike frequency (*n*=93, *p*<0.001), which plateaued at a concentration of 5 mM (1 *vs* 5 mM post-hoc: *n*=68, *p*=0.025, 5 *vs* 10 mM post-hoc: *n*=93, *p*=0.980). In contrast, lysine concentration did not significantly predict spike frequency (figure 2*e*; *n*=65, GLMM: t=-1.07, coefficient estimate -0.022, se 0.021, *p*=0.285). The responses to 5 mM sucrose presented to sensilla subsequently exposed to valine or lysine were not significantly different (*n*=69, Student’s t-test, t(67.0)=0.894, *p*=0.375), indicating that the two groups of bees did not differ in their responses to sugars.

We compared the responses to valine (0.1-50 mM) and sucrose (1-50 mM, figure 2*f*) from sensilla exposed to both tastants. There was a significant interaction between the tastant (sucrose or valine) and concentration (*n*=34, GLMM: t=-3.38, coefficient estimate -0.212 for valine, se 0.063, *p*<0.001): increasing valine concentrations evoked lower spike frequencies than increasing sucrose concentrations (figure 2*g*). The sigmoid fitted to the frequency response, *f*, of GRNs to increasing sucrose concentrations, *c*, was *f*=32.1/(1+e^-(c-3.91)0.399^). The sigmoid fitted to the frequency response of GRNs stimulated with increasing valine concentrations was *f*=20.5/(1+e^-(c-0.699)1.34^). Sucrose had a higher frequency plateau (*f*_max_=32.1 Hz) and half-maximal effective concentration (EC_50_=3.91 mM) than valine (*f*_max_=20.5 Hz, EC_50_=0.699 mM). Thus, gustatory sensilla responded to valine with a lower frequency plateau, reached at lower concentrations, compared to sucrose.

## 4. Discussion

We demonstrate that bumblebees can perceive the EAA valine pre-ingestively via gustatory sensilla on their mouthparts, specifically the galea. This response was dose-dependent, reaching an asymptote at 5 mM. In contrast, the response of sensilla to the EAA lysine was not significantly different from water, suggesting bumblebees are unable to perceive lysine pre-ingestively via galeal A-type gustatory sensilla between 0.1 and 500 mM. An increase in spike rate was first elicited for 0.5 mM valine, and the EC_50_ (half maximal effective concentration) was 0.7 mM, suggesting that bumblebees are sensitive to lower concentrations of valine than sucrose (EC_50_ = 3.91 mM). The absence of burst-like firing in response to valine, even at high concentrations, suggests differing mechanisms for valine and sucrose detection, as burst firing has been previously hypothesized to prevent sensory adaptation to high sucrose concentrations [27]. Thus, bumblebees’ GRNs are both specific and sensitive in their responses to at least one EAA, valine.

Other studies have also observed heterogenous responses of bees to amino acids, at the molecular, electrophysiological and behavioural level [20,21,24,26]. Our results contrast with previous behavioural observations. When amino acids were presented to the antennae in a differential proboscis extension response (PER) assay, bumblebees could not distinguish valine (85 mM) from water but could distinguish lysine (8.5-170 mM) within the concentration ranges we used to stimulate sensilla. While the PER method does not distinguish between olfactory, tactile and gustatory cues, which might also contribute to amino acid perception, such a discrepancy could also arise from differential expression of receptors between the antenna and mouthparts [31] or from age and/or experience-related differences in receptor expression between the bees tested in the two studies [32]. We selected bumblebee foragers observed collecting pollen, whereas Ruedenauer et al. [24] selected workers, not specifying whether they were foraging or in-hive bees.

One other study has performed single sensillum recordings from honeybees (*Apis mellifera*) to measure the sensitivity of bee mouthparts to NEAAs [26], finding a dose-dependent response to glutamate and aspartate. Sensilla responded to these NEAAs with an increasing response between 50-200 mM. Voltage-clamp recordings and calcium imaging of genetically expressed *AmGr10*, a gustatory receptor (GR) also present in *Bombus terrestris*, showed a response to multiple amino acids to differing degrees, but not to sweet or bitter substances. Expressed *AmGr10* exhibited sensitivity to lysine but not valine, again contrasting with our findings from bumblebee mouthpart recordings. While *AmGr10* is expressed in the honeybee mouthparts, there is much higher expression in the antennae [26], which, as well as potential species-specific differences in receptor types and expression, could account for the discrepancy in findings. Interestingly, Lim *et al*. [26] also observed that sensitivity to both lysine and valine could be enhanced by the addition of purine ribonucleotides that are also found in pollen [26] and may therefore play a role in enhancing nutrient perception by bees.

Much of our understanding of gustatory perception in insects, particularly beyond behavioural assays, derives from studies on flies. Their gustatory responses to amino acids are also compound- and concentration-dependent [33,34] and vary according to sensilla type [35]. Several mechanisms are thought to be involved in amino acid perception at the mouthparts, involving both sweet-sensing [35– 37] and bitter-sensing [35] GRNs. Furthermore, amino acid perception relies on both GRs and ionotropic receptors (IRs) [34,35]. *Drosophila melanogaster* labellar cells show a biphasic response to some amino acids: lysine elicits a weakly attractive response via the sweet-sensing GRNs at low concentrations, whereas high concentrations are aversive and activate the IRs in bitter-sensing GRNs [34]. Valine in contrast, elicited a response in bitter-sensing GRNs at both low and high concentrations. For both valine and lysine perception, IRs were required. The *Drosophila melanogaster* genome contains 66 IR genes, whereas the *Bombus terrestris* genome contains just 21 IR genes [38], though no studies that have examined their function in bee gustation. Some flies express valine-gated IRs in their labellar sweet taste cells [36], raising the question of whether valine is similarly detected via sugar-sensitive GRNs or a dedicated GRN sensitive to some, but not all, amino acids. Combined with the absence of lysine response in bumblebee mouthpart sensilla, this suggests further work is needed to conclude whether bumblebees truly possess dedicated amino acid receptors, as is suspected of honeybees [26].

This newly demonstrated capacity for pre-ingestive amino acid perception provides a potential mechanism for bumblebees to rapidly assess floral rewards while foraging based on amino acid as well as sugar content, which may guide their foraging choices and have significant implications for bee-flower interactions and pollination. Bumblebees responded to valine at 0.5 mM, meaning they may be able to perceive it in nectar, where concentrations rarely exceed 1 mM [12–15], as well as pollen, where amino acids are found, both free and bound in proteins, between 10-200 mM [8–11]. It is already known that bees have dose-dependent feeding preferences for amino acids when presented in sugar water, and so it is likely that amino acid concentrations, which vary considerably between the nectar and pollen of different plant species or even between plants within the same species [6,7], could impact individual foraging choices in real-time, without the need for post-ingestive cues or feedback from nestmates on return to the colony [39]. While the mouthparts of bumblebees undoubtedly come into contact with pollen during flower handling and the addition of nectar for pollen packing into the corbiculae [40], there remain open questions regarding the opportunities for adult bees to sample nutritional cues from pollen pre-ingestively, since the majority of free amino acids are found inside the pollen grain [5]. Evidence from other insects, such as butterflies, shows that pollen grains can be lysed at the mouthparts through osmotic and mechanical processes to release the nutritional contents inside [39,41]. If these mechanisms also occur at the bumblebee mouthparts, then bees would also be able to detect the presence and concentration of certain amino acids during pollen collection. Further work is needed to determine whether bees evaluate the nutritional quality of pollen via the presence or specific concentration of a single common amino acid or rather use a combination of amino acids (*i*.*e*. those most found across different pollen species) as a more reliable gustatory cue.

## Author contributions

SR, JEN and EN designed the study. SR conducted the electrophysiological recordings and statistical analysis. RHP provided advice regarding the set-up, data extraction and statistical analysis. SR and EN wrote the first draft of the manuscript. All authors contributed to and approve of the final draft.

## Competing interests

We declare we have no competing interests.

## Funding

This work was funded by a UKRI Future Leaders Fellowship awarded to EN (MR/T021691/1).

## Acknowledgements

We are grateful to Geraldine Wright, Ashwin Miriyala, and Sooho Lim for helpful advice regarding electrophysiological recordings and tastant preparation.

## Notes

### Competing Interest Statement

The authors have declared no competing interest.

### Summary of Updates

Minor changes to streamline publication process

## References

1. Heyneman AJ. 1983 Optimal sugar concentrations of floral nectars - dependence on sugar intake efficiency and foraging costs. Oecologia 60, 198–213. (doi:10.1007/BF00379522)

2. Núñez JA. 1970 The relationship between sugar flow and foraging and recruiting behaviour of honey bees (Apis mellifera L.). Anim. Behav. 18, 527–538. (doi:10.1016/0003-3472(70)90049-7)

3. Lihoreau M, Chittka L, Raine NE, Kudo G. 2011 Trade-off between travel distance and prioritization of high-reward sites in traplining bumblebees. Funct. Ecol. 25, 1284–1292. (doi:10.1111/j.1365-2435.2011.01881.x)

4. Wright GA, Nicolson SW, Shafir S. 2018 Nutritional physiology and ecology of honey bees. Annu. Rev. Entomol. 63, 327–344. (doi:10.1146/annurev-ento-020117-043423)

5. Nicolson SW. 2011 Bee food: the chemistry and nutritional value of nectar, pollen and mixtures of the two. African Zool. 46, 197–204. (doi:10.1080/15627020.2011.11407495)

6. Gottsberger G, Schrauwen J, Linskens HF. 1984 Amino acids and sugars in nectar, and their putative evolutionary significance. Plant Syst. Evol. 145, 55–77. (doi:10.1007/BF00984031)

7. Gardener MC, Gillman MP. 2001 Analyzing variability in nectar amino acids: composition is less variable than concentration. J. Chem. Ecol.27, 2545–2558. (doi:10.1023/A:1013687701120)

8. Jeannerod L, Carlier A, Schatz B, Daise C, Richel A, Agnan Y, Baude M, Jacquemart A-L. 2022 Some bee-pollinated plants provide nutritionally incomplete pollen amino acid resources to their pollinators. PLoS One 17, e0269992. (doi:10.1371/journal.pone.0269992)

9. Weiner CN, Hilpert A, Werner M, Linsenmair KE, Blüthgen N. 2010 Pollen amino acids and flower specialisation in solitary bees. Apidologie41, 476–487. (doi:10.1051/apido/2009083)

10. Stabler D, Power EF, Borland AM, Barnes JD, Wright GA. 2017 A method for analysing small samples of floral pollen for free and protein-bound amino acids. Methods Ecol. Evol. 9, 430–438. (doi:10.1111/2041-210X.12867)

11. Szczesna T. 2006 Protein content and amino acids composition of bee-collected pollen originating from Poland, South Korea and China. J. Apic. Sci. 50, 91–99.

12. Power EF, Stabler D, Borland AM, Barnes J, Wright GA. 2018 Analysis of nectar from low-volume flowers: A comparison of collection methods for free amino acids. Methods Ecol. Evol. 9, 734–743. (doi:10.1111/2041-210X.12928)

13. Baker HG, Baker I. 1986 The occurrence and significance of amino acids in floral nectar. Plant Syst. Evol. 151, 175–186. (doi:10.1007/BF02430273)

14. Baker HG, Baker I. 1973 Amino-acids in nectar and their evolutionary significance. Nature 241, 543–545. (doi:10.1038/241543b0)

15. Parkinson RH, Power EF, Walter K, McDermott-Roberts AE, Pattrick JG, Wright GA. 2024 Do pollinators play a role in shaping the essential amino acids found in nectar? bioRxiv., 2024.09.30.615756. (doi:10.1101/2024.09.30.615756)

16. Inouye DW, Waller GD. 1984 Responses of honey bees (Apis mellifera) to amino acid solutions mimicking floral nectars. Ecology 65, 618–625. (doi:10.2307/1941424)

17. Alm J, Ohnmeiss TE, Lanza J, Vriesenga L. 1990 Preference of cabbage white butterflies and honey bees for nectar that contains amino acids. Oecologia 84, 53–57. (doi:10.1007/BF00665594)

18. Kim YS, Smith BH. 2000 Effect of an amino acid on feeding preferences and learning behavior in the honey bee, Apis mellifera. J. Insect Physiol. 46, 793–801. (doi:10.1016/S0022-1910(99)00168-7)

19. Cook DL, Schwindt PC, Grande LA, Spain WJ. 2003 Synaptic depression in the localization of sound. Nature 421, 66–70. (doi:10.1038/nature01561)

20. Simcock NK, Gray HE, Wright GA. 2014 Single amino acids in sucrose rewards modulate feeding and associative learning in the honeybee. J. Insect Physiol. 69, 41–48. (doi:10.1016/J.JINSPHYS.2014.05.004)

21. Carter C, Shafir S, Yehonatan L, Palmer RG, Thornburg R. 2006 A novel role for proline in plant floral nectars. Naturwissenschaften 93, 72–79. (doi:10.1007/s00114-005-0062-1)

22. de Groot AP. 1953 Protein and Amino Acid Requirements of the Honeybee (Apis Mellifica L.).

23. Hendriksma HP, Shafir S. 2016 Honey bee foragers balance colony nutritional deficiencies. Behav. Ecol. Sociobiol. 70, 509–517. (doi:10.1007/s00265-016-2067-5)

24. Ruedenauer FA, Leonhardt SD, Lunau K, Spaethe J. 2019 Bumblebees are able to perceive amino acids via chemotactile antennal stimulation. J. Comp. Physiol. A 205, 321–331. (doi:10.1007/S00359-019-01321-9)

25. Linander N, Hempel de Ibarra N, Laska M. 2012 Olfactory detectability of L-amino acids in the European honeybee (Apis mellifera). Chem. Senses 37, 631–638. (doi:10.1093/chemse/bjs044)

26. Lim S, Jung J, Yunusbaev U, Ilyasov R, Kwon HW. 2019 Characterization and its implication of a novel taste receptor detecting nutrients in the honey bee, Apis mellifera. Sci. Rep. 9, 11620. (doi:10.1038/s41598-019-46738-z)

27. Miriyala A, Kessler S, Rind FC, Wright GA. 2018 Burst firing in bee gustatory neurons prevents adaptation. Curr. Biol. 28, 1585–1594.e3. (doi:10.1016/J.CUB.2018.03.070)

28. Marion-Poll F, van der Pers J. 1996 Un-filtered recordings from insect taste sensilla. Entomol. Exp. Appl. 80, 113–115. (doi:10.1111/j.1570-7458.1996.tb00899.x)

29. Parkinson RH, Kessler SC, Scott J, Simpson A, Bu J, Al-Esawy M, Mahdi A, Miriyala A, Wright GA. 2022 Temporal responses of bumblebee gustatory neurons to sugars. iScience 25, 104499. (doi:10.1016/j.isci.2022.104499)

30. R Core Team. 2013 R: A language and environment for statistical computing.

31. Simcock NK, Wakeling LA, Ford D, Wright GA. 2017 Effects of age and nutritional state on the expression of gustatory receptors in the honeybee (Apis mellifera). PLoS One 12, e0175158. (doi:10.1371/journal.pone.0175158)

32. Claudianos C, Lim J, Young M, Yan S, Cristino AS, Newcomb RD, Gunasekaran N, Reinhard J. 2014 Odor memories regulate olfactory receptor expression in the sensory periphery. Eur. J. Neurosci. 39, 1642–54. (doi:10.1111/ejn.12539)

33. Park J, Carlson JR. 2018 Physiological responses of the Drosophila labellum to amino acids. J. Neurogenet. 32, 27–36. (doi:10.1080/01677063.2017.1406934)

34. Ganguly A, Pang L, Duong V-K, Lee A, Schoniger H, Varady E, Dahanukar A. 2017 A molecular and cellular context-dependent role for Ir76b in detection of amino acid taste. Cell Rep. 18, 737–750. (doi:10.1016/j.celrep.2016.12.071)

35. Aryal B, Dhakal S, Shrestha B, Lee Y. 2022 Molecular and neuronal mechanisms for amino acid taste perception in the Drosophila labellum. Curr. Biol. 32, 1376–1386.e4. (doi:10.1016/j.cub.2022.01.060)

36. Murakami M, Kijima H. 2000 Transduction ion channels directly gated by sugars on the insect taste cell. J. Gen. Physiol. 115, 455–466. (doi:10.1085/jgp.115.4.455)

37. Shimada I. 1978 The stimulating effect of fatty acids and amino acid derivatives on the labellar sugar receptor of the fleshfly. J. Gen. Physiol.71, 19–36. (doi:10.1085/jgp.71.1.19)

38. Sadd BM et al. 2015 The genomes of two key bumblebee species with primitive eusocial organization. Genome Biol. 16, 76. (doi:10.1186/s13059-015-0623-3)

39. Pernal S, Currie R. 2001 The influence of pollen quality on foraging behavior in honeybees (Apis mellifera L.). Behav. Ecol. Sociobiol. 51, 53–68. (doi:10.1007/s002650100412)

40. Woodrow C, Jafferis N, Kang Y, Vallejo-Marín M. 2024 Buzz-pollinating bees deliver thoracic vibrations to flowers through periodic biting. Curr. Biol. 34, 4104–4113.e3. (doi:10.1016/j.cub.2024.07.044)

41. Krenn HW, Eberhard MJB, Eberhard SH, Hikl A-L, Huber W, Gilbert LE. 2009 Mechanical damage to pollen aids nutrient acquisition in Heliconius butterflies (Nymphalidae). Arthropod. Plant. Interact. 3, 203–208. (doi:10.1007/s11829-009-9074-7)

